# Neural Population Control via Deep Image Synthesis

**DOI:** 10.1101/461525

**Authors:** Pouya Bashivan, Kohitij Kar, James J DiCarlo

## Abstract

Particular deep artificial neural networks (ANNs) are today’s most accurate models of the primate brain’s ventral visual stream. Here we report that, using a targeted ANN-driven image synthesis method, new luminous power patterns (i.e. images) can be applied to the primate retinae to predictably push the spiking activity of targeted V4 neural sites beyond naturally occurring levels. More importantly, this method, while not yet perfect, already achieves unprecedented independent control of the activity state of entire *populations* of V4 neural sites, even those with overlapping receptive fields. These results show how the knowledge embedded in today’s ANN models might be used to non-invasively set desired internal brain states at neuron-level resolution, and suggest that more accurate ANN models would produce even more accurate control.

---

Particular deep feed-forward artificial neural network models (ANNs) constitute today’s most accurate “understanding” of the initial ~200ms of processing in the primate ventral visual stream and the core object recognition behavior it supports (see (*1*) for the currently leading models). In particular, visually-evoked internal neural representations of these specific ANNs are remarkably similar to the visually-evoked neural representations in mid-level (area V4) and high-level (area IT) cortical stages of the ventral stream (*2, 3*), a finding that has been extended to neural representations in visual area V1 (*4*), to patterns of behavioral performance in core object recognition tasks (*5, 6*), and to both magnetoencephalography and fMRI measurements from the human ventral visual stream (*7, 8*). Notably, these prior findings of model-to-brain similarity were not curve fits to brain data – they were *predictions* evaluated using images not previously seen by the ANN models, showing that these models have some generalization of their ability to capture key functional properties of the ventral visual stream.

However, at least two important potential limitations of this claim have been raised. First, because the visual processing that is executed by the models is not simple to describe, and the models have only been evaluated in terms of internal functional similarity to the brain (above), perhaps they are more like a copy of, rather than a useful “understanding” of, the ventral stream. Second, because the images to assess similarity were sampled from the same distribution as that used to set the model’s internal parameters (photograph and rendered object databases), it is unclear if these models would pass a stronger test of functional similarity – does that similarity generalize to entirely novel images? That is, perhaps their reported apparent functional similarity to the brain (*3, 7, 9*), substantially over-estimates their true functional similarity.

Here we conducted a set of non-human primate visual neurophysiology experiments to assess the first potential limitation by asking if the detailed knowledge that the models contain is useful for one potential application (neural activity control), and to assess the second potential limitation by asking if the functional similarity of the model to the brain generalizes to entirely novel images.

Specifically, we used one of the leading deep ANN ventral stream models (i.e. a specific model with a fully fixed set of parameters) to synthesize new patterns of luminous power (“controller images”) that, when applied to the retinae, were intended to control the neural firing activity of particular, experimenter-chosen neural sites in cortical visual area V4 of macaques in two settings. i) *Neural “Stretch”*: synthesize images that “stretch” the maximal firing rate of any single targeted neural site well beyond its naturally occurring maximal rate. ii) *Neural Population State Control*: synthesize images to independently control every neural site in a small recorded population (here, populations of 5-40 neural sites). We here tested that population control by aiming to use such model-designed retinal inputs to drive the V4 population into an experimenter-chosen “one hot” state in which one neural site is pushed to be highly active while all other nearby sites are simultaneously all “clamped” at their baseline activation level. We reasoned that successful experimenter control would demonstrate that at least one ANN model can be used to non-invasively *control* the brain – a practical test of useful, causal “understanding” (*10, 11*).

We used chronic, implanted microelectrode arrays to record the responses of 107 neural multi-unit and single-unit sites from visual area V4 in three awake, fixating rhesus macaques (*n_M_* =52, *n_N_* =33, *n_S_*=22). We first determined the classical receptive field (cRF) of each site with briefly presented small squares (for details see Methods). We then tested each site using a set of 640 naturalistic images (always presented to cover the central 8° of the visual field that overlapped with the estimated cRFs of all the recorded V4 sites), and using a set of 370 complex curvature stimuli previously determined to be good drivers of V4 neurons (*12*) (location tuned for the cRFs of the neural sites). Using each site’s visually evoked responses (see Methods) to 90% of the naturalistic images (n=576), we created a mapping from a single “V4” layer of a deep ANN model (*13*) (Conv-3 layer; that we had established in prior work) to the neural responses. The predictive accuracy of this model-to-brain mapping has previously been used as a measure of the functional fidelity of the brain model to the brain (*1, 3*). Indeed, using the V4 responses to the held-out 10% of the naturalistic images as tests, we replicated and extended that prior work – we found that the neural predictor models correctly predicted 89% of the explainable (i.e. image driven) variance in the V4 neural responses (median over the 107 sites, each site computed as the mean over two mapping/testing splits of the data; see Methods).

Besides generating a model-V4-to-brain-V4 similarity score (89%, above), this mapping procedure produces a potentially powerful tool – an image-computable predictor model of the visually-evoked firing rate of each of the V4 neural sites. If truly accurate, this predictor model is not simply a data fitting device and not just a similarity scoring method – instead it must implicitly capture a great deal of visual “knowledge” that may be difficult to express in human language, but is hypothesized (by the model) to be used by the brain to achieve successful visual behavior. To extract and deploy that knowledge, we used a model-driven image synthesis algorithm (see Figure-1 and Methods) to generate *controller images* that were customized for each neural site (i.e. according to its predictor model) so that each image should predictably and reproducibly *control* the firing rates of V4 neurons in a particular, experimenter-chosen way. That is, we aimed to test the hypothesis that experimenter-delivered application of a particular pattern of luminous power on the retinae will reliably and reproducibly *cause* V4 neurons to move to a particular, experimenter-specified activity state (and that removal of that pattern of luminous power will return those V4 neurons to their background firing rates).

**Figure 1:**
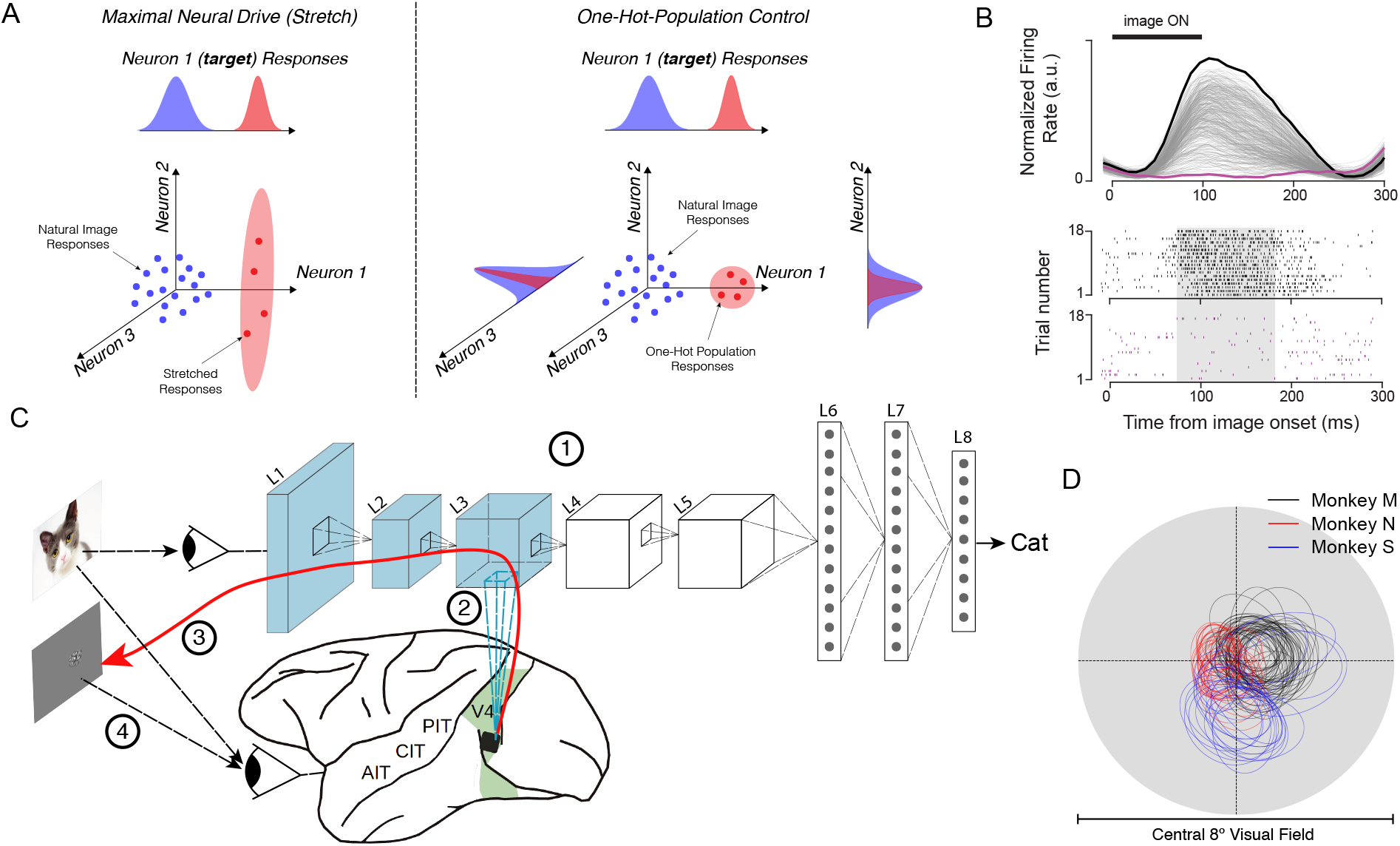
Overview of the synthesis procedure. A) Schematic illustration of the two tested control scenarios. Left - the controller algorithm synthesizes novel images that it believes will maximally drive the firing rate of a target neural site (Stretch). In this case, the controller algorithm does not attempt to regulate the activity of other measured neurons (e.g. they might also increase as shown). Right - the controller algorithm synthesizes images that it believes will maximally drive the firing rate of a target neural site while suppressing the activity of other measured neural sites (one-hot population). B) Top - gray lines (overlapping): responses of a single example V4 neural site to 640 naturalistic images (averaged over 40 repetitions for each image). Vertical wide black line marks the image presentation period. Bottom - raster plots of highest (black) and lowest (purple) neural response to naturalistic images. Shaded area indicates the time window over which the activity level of each V4 neural site is computed (i.e. one value per image for each neural site). C) The neural control experiments are done in four steps. (1) Parameters of the neural network are optimized by training on a large set of labeled natural images (Imagenet (*27*)) and then held constant thereafter. (2) ANN “neurons” are mapped to each recorded V4 neural site. The mapping function constitutes an image-computable predictive model of the activity of each of those V4 sites. (3) The resulting differentiable model is then used to synthesize “controller” images for either single-site or population control. (4) The luminous power patterns specified by these images are then applied by the experimenter to the subject’s retinae and the degree of control of the neural sites is measured. D) Classical receptive fields of neural sites in monkey M (black), Monkey N (red) and Monkey S (blue; see Methods).

While there are an extremely large number of possible neural activity states that an experimenter might ask a controller method to try to achieve, we restricted our experiments to the V4 spiking activity 70-170 ms after retinal power input (the time frame where the ANN models are presumed to be most accurate), and we have thus far tested two control settings: *Stretch control* and *One-hot population control* (described below). To test and quantify the goodness of control, we applied patterns of luminous power specified by the synthesized *controller images* to the retinae of the animal subjects while we recorded the responses of the same V4 neural sites (see Methods).

Each experimental manipulation of the pattern of luminous power on the retinea are colloquially referred to as “presentation of an image”, but we state the precise manipulation here of applied power that is under experimenter control and fully randomized with other applied luminous power patterns (other images) to emphasize that this is logically *identical* to more direct energy application (e.g. optogenetic experiments) in that the goodness of experimental control is inferred from the correlation between power *manipulation* and the neural response in exactly the same way in both cases (see (*11*) for review). The only difference of the two approaches is the assumed mechanisms that intervene between the experimentally-controlled power and the controlled dependent variable (here V4 spiking rate) – steps that the ANN model aims to approximate with stacked synaptic sums, threshold non-linearities, and normalization circuits. In both the control case presented here and the optogenetics control case, those intervening steps are not fully known, but approximated by a model of some type. That is, neither experiment is “only correlational” because causality is inferred from experimenter-delivered, experimenter-randomized application of power to the system.

Because each experiment was performed over separate days of recording (one day to build all the predictor models, one day to test control), only neural sites that maintained both high SNR and consistent rank order of responses to a standard set of 25 naturalistic images across the two experimental days were considered further (*n_M_* =38, *n_N_* =19, and *n_S_*=19 for *Stretch* experiments; *n_M_* =38, and *n_S_*=19 for *One-hot-population* experiments; see Methods).

## “Stretch” Control: Attempt to maximize the activity of individual V4 neural sites

We first defined each V4 site’s “naturally-observed maximal firing rate” as that which was found by testing its response to the best of the 640 naturalistic test images (cross-validated over repeated presentations, see Methods). We then generated synthetic *controller images* for which the synthesis algorithm was instructed to drive one of the neural site’s firing rate as high as possible beyond that rate, regardless of the other V4 neural sites. For our first Stretch Control experiment, we restricted the synthesis algorithm to only operate on parts of the image that were within the classical receptive field (cRF) of each neural site. For each target neural site (*n_M_* =21, *n_N_* =19, and *n_S_*=19), we ran the synthesis algorithm from five different random image initializations. For 79% of neural sites, the synthesis algorithm successfully found at least one image that it predicted to be at least 10% above the site’s naturally observed maximal firing rate (see Methods). However, in the interest of presenting an unbiased estimate of the stretch control goodness for randomly sampled V4 neural sites, we included all sites in our analyses, even those (~20%) that the control algorithm predicted that it could not ”stretch.” Visual inspection suggests that the five *stretch controller* images generated by the algorithm for each neural site are perceptually more similar to each other compared to those generated for different neural site (see Figures 2 and S1), but we did not psychophysically quantify that similarity.

**Figure 2:**
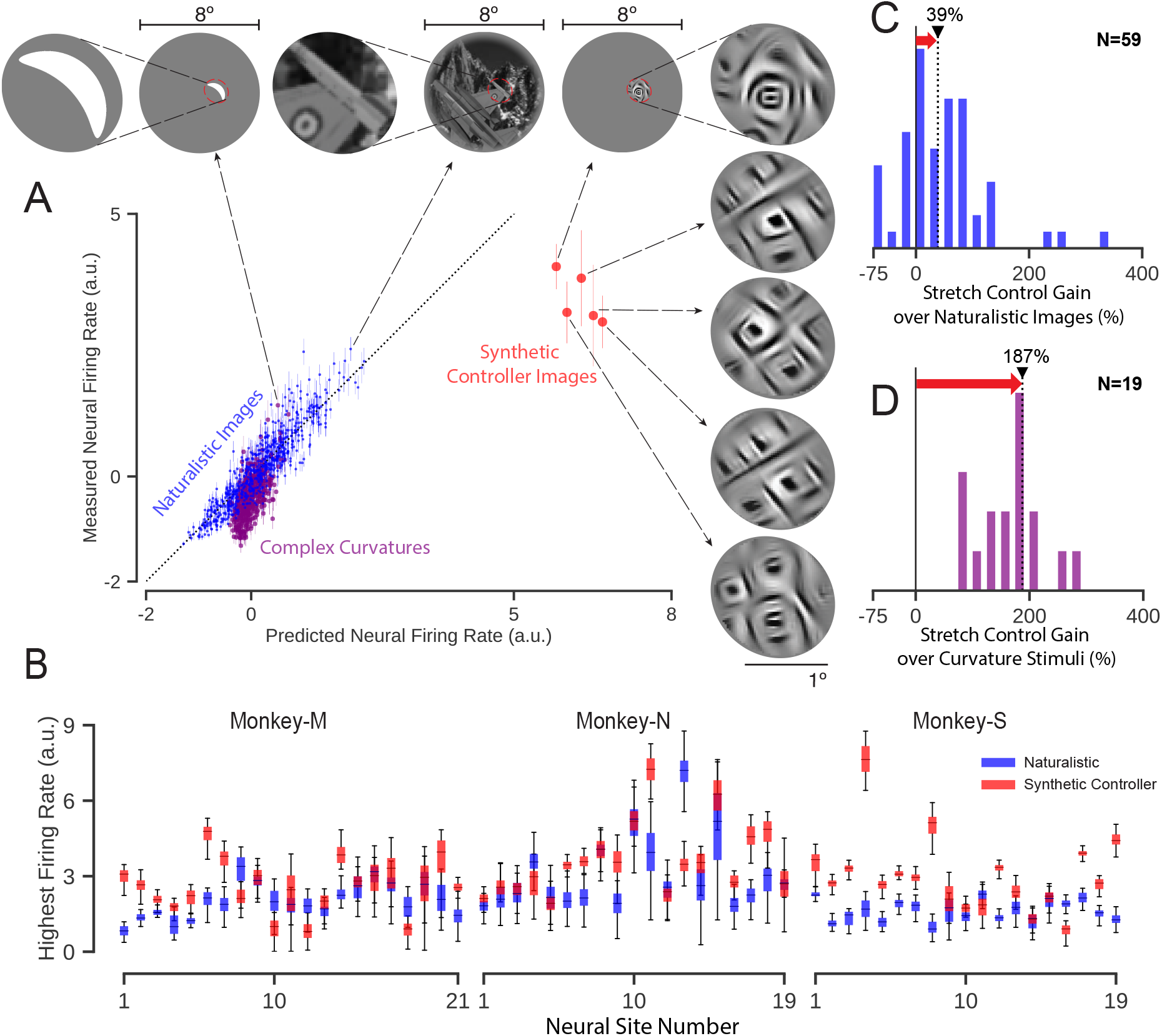
Maximal drive of individual neural sites (Stretch). A) Results for an example successful “stretch” control test. Normalized activity level of the target V4 neural sites is shown for all of the naturalistic images (blue dots), complex curved stimuli (purple dots) and for its five synthetic “stretch” controller images (red dots; see Methods). Best driving images within each category, and the zoomed view of the receptive field are shown on the top. B) Difference in firing rate in response to naturalistic (blue boxes) and synthetic images (red boxes) for each neural site in three monkeys. Controller image synthesis was restricted within the receptive field of the target neural site. C) Histogram of increase in the firing rate over naturalistic images for cRF-restricted synthetic images. D) Histogram of increase in the firing rate over complex curved stimuli. Black triangle with dotted black line marks the median of the scores over all tested neural sites. The red arrow highlights the gain in firing rate in each experiment achieved by the controller images. “N” indicates the number of neural sites included in each experiment.

An example of the results of applying the *Stretch Control* images to the retinae of one monkey to target one of its V4 sites is shown in Figure 2-A), along with the ANN-model-predicted responses of this site for all tested images. A closer visual inspection of this neural site’s “best” natural and complex curvature images within the site’s cRF (Fig. 2 top) suggests that it might be especially sensitive to the presence of an angled convex curvature in the middle and a set of concentric circles at the bottom left side. This is consistent with extensive systematic work in V4 using such stimuli (*12, 14*), and it suggests that we had successfully located the cRF and tuned our stimulus presentation to maximize firing rate by the standards of such prior work. Interestingly however, we found that all five synthetic stretch control images (red) drove the neural responses above the response to each and every tested naturalistic image (blue) and above the response to each and every complex curvature stimulus presented within the cRF (purple), (Fig. 2-A).

To quantify the goodness of this stretch control, we measured the neural response to the best of the five synthetic images (again, cross-validated over repeated presentations, see Methods) and compared it with the naturally-observed maximal firing rate (defined above). We found that the stretch controller images successfully drove 68% of the V4 neural sites (40 out of 59) statistically beyond its maximal naturally-observed firing rate (unpaired-samples t-test at the level of *p* < 0.01 between distribution of highest firing rates for naturalistic and synthetic images; distribution generated from 50 random cross-validation samples, see Methods). Measured as an amplitude, we found that the stretch controller images typically produced a firing rate that was 39% higher than the maximal naturalistic firing rate (median over all tested sites, Figure-2 panel B and C).

Because our fixed set of naturalistic images was not optimized to maximally drive each V4 neural site, we considered the possibility that our stretch controller was simply rediscovering image pixel arrangements that are already known from prior systematic work to be good drivers of V4 neurons (*12, 14*). To test this hypothesis, we tested 19 of the V4 sites (*n_M_* = 11, *n_S_* = 8) by presenting – inside the cRF of each neural site – each of 370 complex curve shapes (*14*) – a stimulus set that has been previously shown to contain image features that are good at driving V4 neurons when placed within the cRF. Because we were also concerned that the fixed set of naturalistic images did not maximize the local image contrast within each V4 neuron’s cRF, we presented the complex curved shapes at a contrast that was matched to the contrast of the synthetic stretch controller images (see supplementary Figure S4). Interestingly, we found that for each tested neural site, the synthetic controller images generated higher firing rates than the most-effective complex curve shape (Figure 2-D). Specifically, when we used the maximal response over all the complex curve shapes as the reference (again, cross-validated over repeated presentations), we found that the median stretch amplitude was even larger (187%) than when the maximal naturalistic image was used as the reference (73% for the same 19 sites). In sum, the ANN-driven stretch controller had discovered pixel arrangements that were better drivers of V4 neural sites than prior systematic attempts to do so.

## “One-Hot-Population” Control: Attempt to *only* activate one of many V4 neural sites

Similar to prior single unit visual neurophysiology studies (*15–17*), the stretch control experiment attempted to optimize the response of each V4 neural site one at a time without regard to the rest of the neural population. But the ANN model potentially enables much richer forms of *population* control in which each neural site might be *independently* controlled. As a first test of this, we asked the synthesis algorithm to try to generate controller images with the goal of driving the response of only one “target” neural site high while *simultaneously* keeping the responses of all other recorded neural sites low (aka a “one-hot” population activity state; see Methods).

We attempted this one-hot-population control on neural populations in which all sites were simultaneously recorded (*One-hot-population* Experiment 1; n=38 in monkey-M; Experiment 2; n=19 in monkey-S). Specifically, we randomly chose a subset of neural sites as “target” sites (14 in monkey-M and 19 in monkey-S) and we asked the synthesis algorithm to generate five one-hot-population controller images for each of these sites (i.e. 33 tests in which each test is an attempt to maximize the activity of one site while suppressing the activity of all other measured sites from the same monkey). For these control tests, we allowed the controller algorithm to optimize pixels over the entire 8° diameter image (that included the cRFs of all the recorded neural sites, see Fig. 3), and we then applied the one-hot-population controller images to the monkey retinea to assess the goodness of control. The synthesis procedure predicted a softmax score of at least 0.5 for 77% of population experiments (as a reference, the maximum softmax score is 1 and is obtained when only the target neural site is active and all off-target neural sites are completely inactive; for an example near 0.3 see Fig. 3).

**Figure 3:**
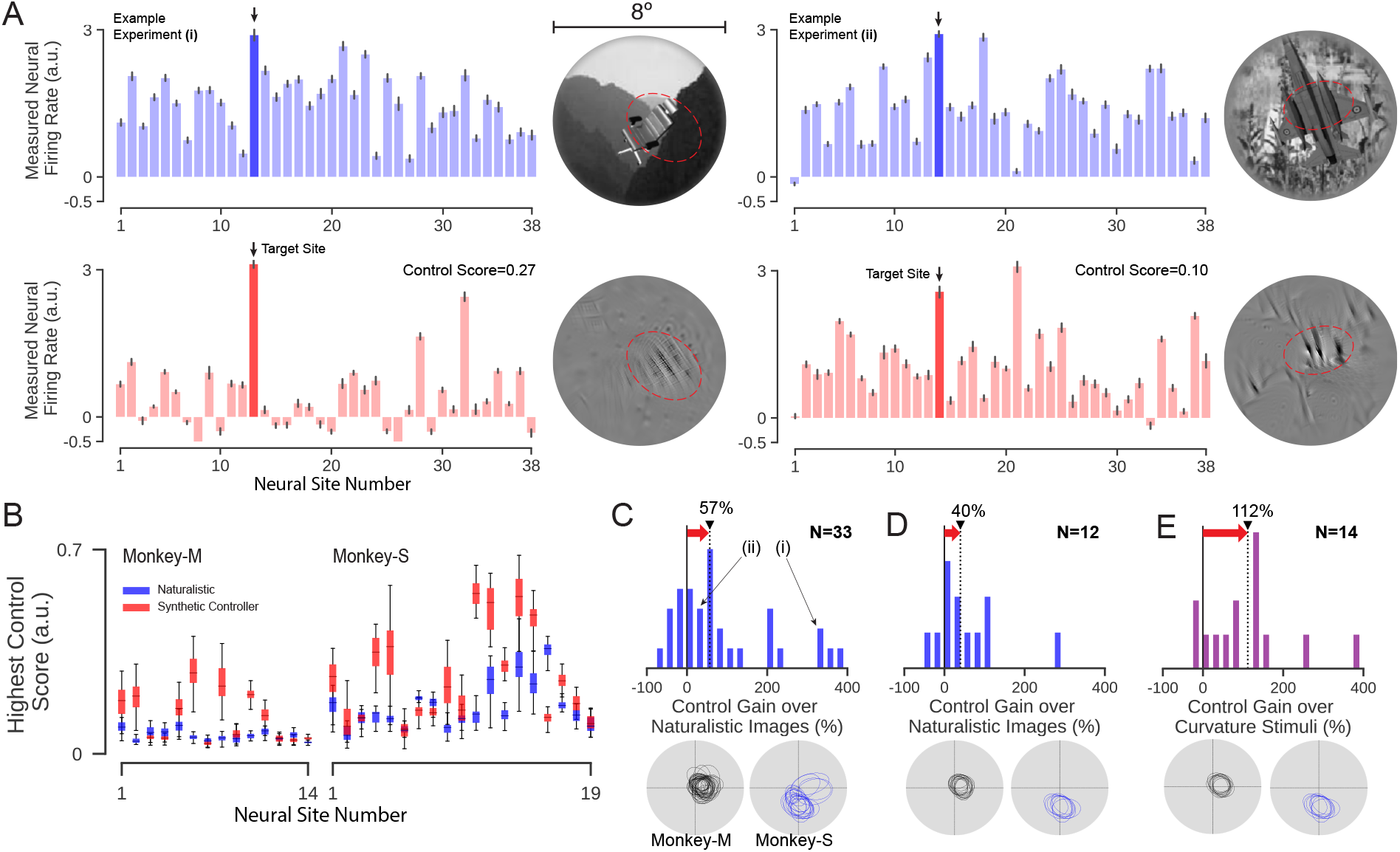
Neural Population Control. We synthesized controller images that aimed to set the neural population in a one-hot state (OHP) in which one target neural site is active and all other recorded neural sites are suppressed. A) Two example OHP experiments (left and right). In each case, the neural activity of each of the validated V4 sites (see Methods) in the recorded population are plotted (most have overlapping cRFs), with the target V4 site indicated in dark blue/red). Note that responses are normalized individually on a normalizer image set to make side-by-side comparison of the responses meaningful (see Methods). Upper panel: activity pattern for the best (“best” in the sense of OHP control, see Methods) naturalistic image (shown on the right). Lower panel: activity pattern produced by retinal application of the ANN-model-synthesized controller image (shown on the right). The red dashed line marks the extended receptive field (2-std) of each site. B) Distribution of control scores for best synthetic and naturalistic images for all 33 OHP full-image controller experiments (*n_M_* = 14*, n_S_* = 19). Control Scores are computed using cross-validation (see Methods). C) Histogram of OHP control gain (i.e. improvement over naturalistic images) for results in (B). (i) and (ii) indicate the scores corresponding to example experiments shown in (A). D) Same experimental data as (C) except analyzed for sub-populations selected so that *all* sites have highly overlapping cRFs (see cRFs below). E) OHP control gain where gain is relative to best complex curvature stimulus in the shared cRF (see text) and controller algorithm is also restricted to operate only in that shared cRF (n=14 OHP experiments). Receptive fields of neural sites in each setting (C-E) (black: monkey-M; blue: monkey-S). “N” indicates the number of experiments in each setting. Red arrow highlights the median gain in control (black triangle) achieved in each case.

While the one-hot-population controller images did not achieve perfect one-hot-population control, we found that the controller images were typically able to achieve enhancements in the activity of the target site without generating much increase in off-target sites (relative to naturalistic images; see examples in Figure 3-A). To quantify the goodness of one-hot-population control in each of the 33 tests, we computed a one-hot-population score on the responses of the activity profile of each population (softmax score, see Methods), and we referenced that score to the one-hot-population control score that could be achieved using only the naturalistic images (i.e without the benefit of the ANN model and synthesis algorithm). We took the ratio of those two scores as the measure of improved one-hot population control, and we found that the controller typically achieved an improvement of 57% (median over all 33 one-hot-population control tests; see Fig. 3-B and C) and we found that that improved control was statistically significant for 76% of the one-hot population control tests (25 out of 33 tests; unpaired-samples t-test at the level of *p <* 0.01).

We considered the possibility that the improved population control was resulting from the non-overlapping cRFs that would allow neural sites to be independently controlled simply by restricting image contrast energy to each site’s cRF. To test this possibility, we analyzed a sub-sample of the measured neural population in which all sites had strongly overlapping cRFs (see Fig. 3-D). We considered a neural population of size 10 in monkey-M and of size 8 in monkey-S for this experiment with largely overlapping cRFs (see Fig. 3-D). In total we performed the experiment on 12 target neural sites in two monkeys (4 in monkey-M and 8 in monkey-S) and found that the amplitude of improved control was still 40%. Thus, a large portion of the improved control is the result of specific spatial arrangements of luminous power within the retinal input region shared by multiple V4 neural sites that the ANN-model has implicitly captured and predicted and the synthesis algorithm has successfully recovered (Fig. 4).

**Figure 4:**
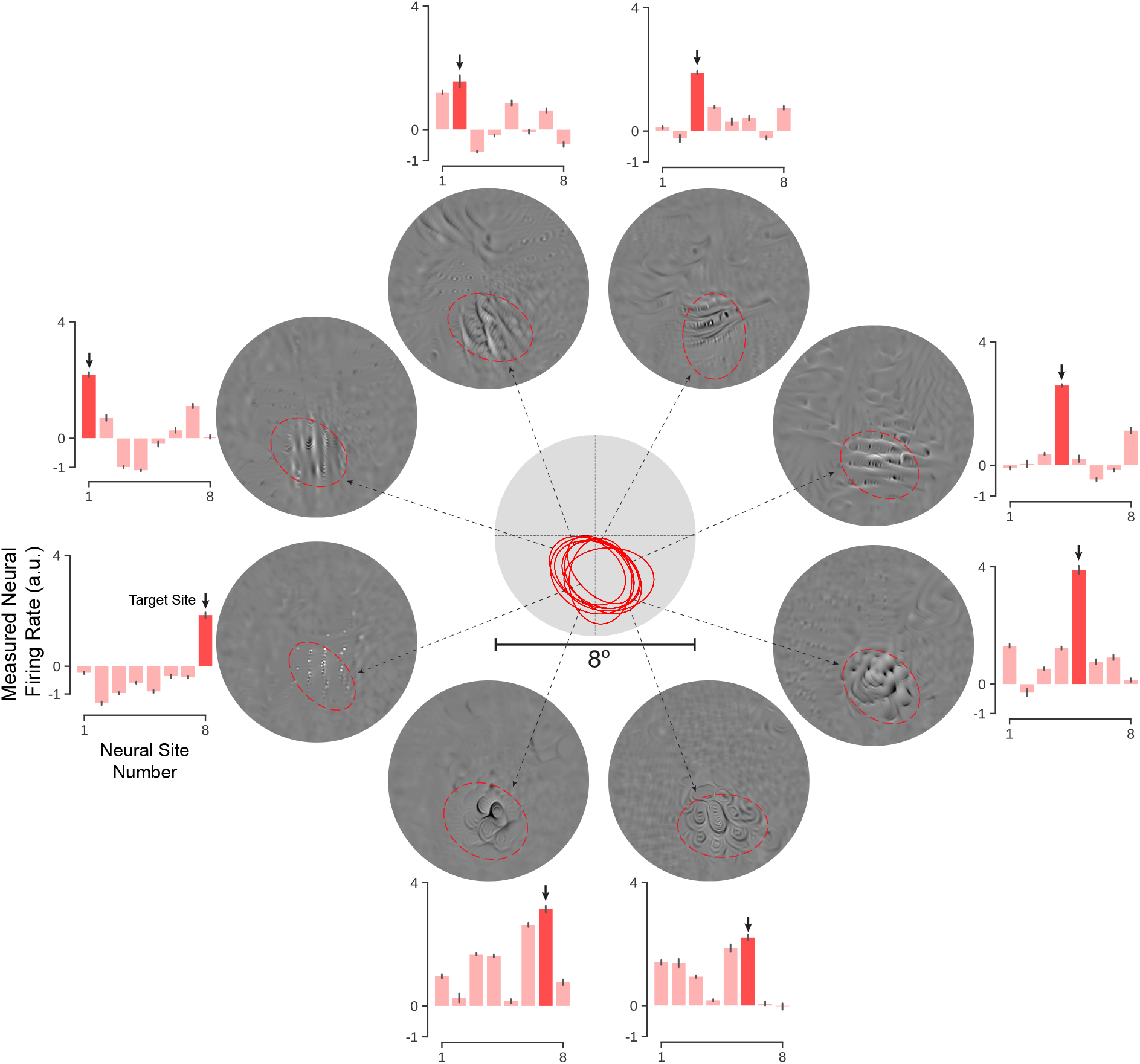
Example of independent control of each neural site on a subset of V4 neural sites with highly overlapping cRFs. Controller images were synthesized to try to achieve a one-hot-population over a population of eight neural sites (in each control test, the target neural site is shown as dark red). Despite highly overlapping receptive fields (center), most of the neural sites could be individually controlled to a reasonable degree. Controller images are shown along with the extended cRF (2-std) of each site (red).

As another test of one-hot-population control, we conducted an additional set of experiments in which we restricted the one-hot control synthesis algorithm to operate only on image pixels within the shared cRF of all neural sites in a sub-population with overlapping cRFs (Fig. 3-E). We compared this within-cRF synthetic one-hot population control with the within-cRF one-hot population control that could be achieved with the complex curved shapes (because the prior experiments with these stimuli were also designed to manipulate V4 responses only using pixels inside the cRF). We found that, for the same set of neural sites, the synthetic controller images produced a very large one-hot population control gain (median 112%, Fig. 3-E) and the control score was significantly higher than best curvature stimulus for 86% of the neural sites (12 out of 14).

## Does the functional fidelity of the ANN brain model generalize to novel images?

Besides testing non-invasive causal neural control, these experiments also aimed to ask if ANN models would pass a stronger test of functional similarity to the brain than prior work had shown (*2, 3*). Specifically, does that model-to-brain similarity generalize to entirely novel images? Because the controller images were synthesized *de novo* from random pixel arrangement and they were optimized to drive the firing rates of V4 neural sites both upwards (targets) and downwards (one-hot-population off-targets), we considered them to be a highly novel set of neural-modulating images that is far removed from the object naturalistic images. Indeed, visual inspection suggests the novelty of these images (Fig. 5). We thus used the V4 neural responses to all the tested synthetic images to ask if the ANN model “neural” responses matched the brain’s responses, using the same similarity measure as prior work (*2, 3*), but now with *zero* parameters to fit. That is, a good model-to-brain similarity score required that the ANN predictor model for each V4 neural site accurately predict the response of that neural site for all of many synthetic images that are each very different than those that we used to train the ANN (photographs) and also very different from the images used to map ANN “V4” sites to individual V4 neural sites (naturalistic images).

**Figure 5:**
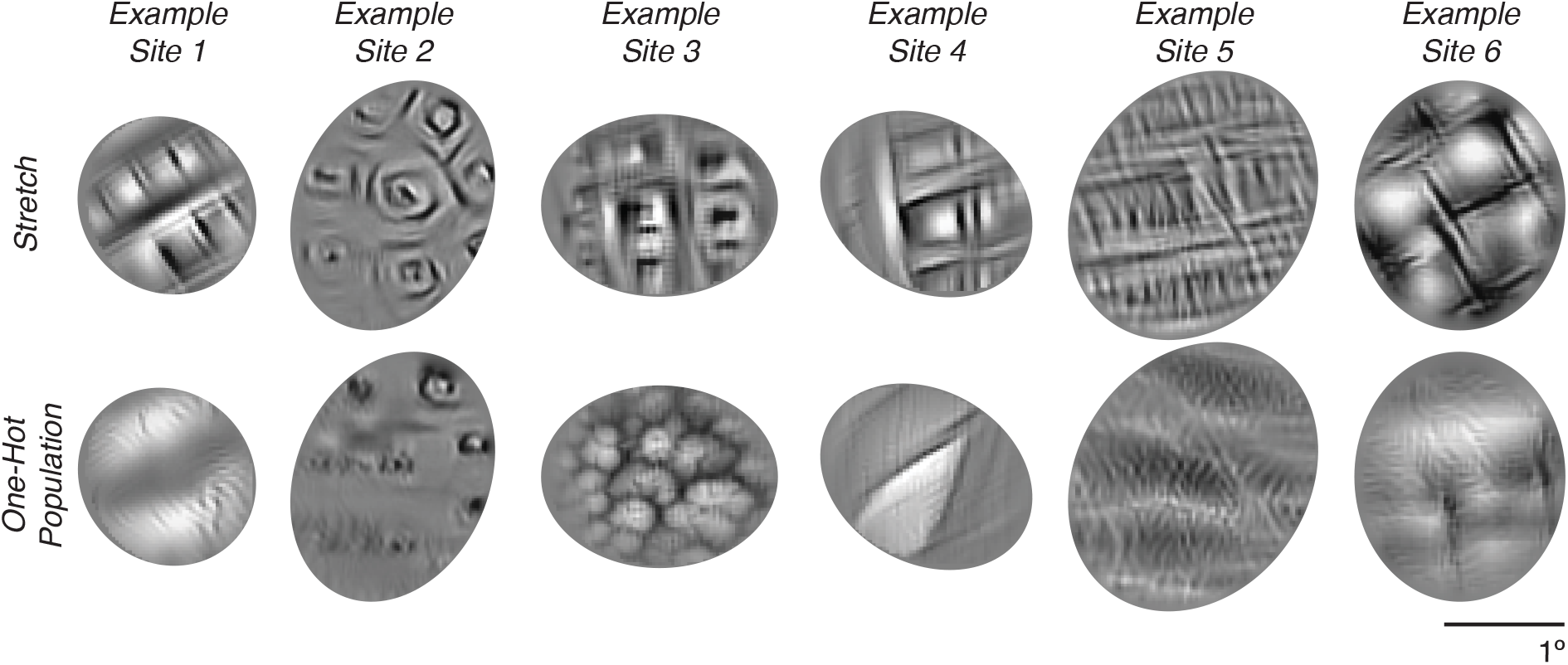
Example controller images synthesized in “Stretch” and “One-hot population” settings for six example target neural sites. Controller images were synthesized from the same initial random image, but optimized for each target neural site and for each control goal (“Stretch” or “One-hot population”, see text). Visual inspection suggests that, for each target site, the One-hot population control images contain only some aspects of the image features in the “Stretch” images.

Consistent with the control results (above), we found that the ANN model accounted for 54% of the explainable variance for the set of synthetic images (median over 76 neural sites in three monkeys; Fig. S3). While this model-to-brain similarity score is lower than that obtained for naturalistic images responses (89%), it is still a substantial portion of the variance considering the fact that all parameters were fixed to make these “out-of-domain” image predictions. We believe this is the strongest test of generalization of today’s ANN models of the ventral stream thus far, and it again shows that the model’s internal neural representation is both remarkably similar to the brain’s intermediate ventral stream representation (V4), but also that it is still not a perfect model of the representation.

## How do we interpret these results?

In sum, we here demonstrate that, using a deep ANN-driven controller method, we can push the firing rates of most V4 neural sites beyond naturally occurring levels and that V4 neural sites with overlapping receptive fields can be partly – but not yet perfectly – independently controlled. In both cases, we show that the goodness of this control is unprecedented in that it is superior to that which can be obtained without the ANN. Finally, we find that – with no parameter tuning at all – the ANN model generalizes quite well to predict V4 responses to synthetic images – images which are strikingly different than the real-world photographs used to tune the ANN synaptic connectivity and map the ANN’s “V4” to each V4 neural site. We believe that these results are the strongest test thus far of today’s deep ANN models of the ventral stream.

Beginning with the work of Hubel and Wiesel (*18, 19*), decades of visual neuroscience has closely equated an understanding of how the brain represents the external visual world with an understanding of what stimuli cause each neuron to respond the most. Indeed, textbooks and important recent results tell us that V1 neurons are tuned to oriented bars (*19*), V2 neurons are tuned to correlated combinations of V1 neurons found in natural images (*20*), V4 neurons are tuned to complex curvature shapes in both 2D and 3D (*16, 21*) and tuned to boundary information (*12, 14*), and IT neurons respond to complex object-like patterns (*17*) including faces (*22, 23*) and bodies as special cases (*24*).

While these efforts have been essential to building both a solid foundation and intuitions about the role of neurons in encoding visual information, our results here show how they can be further refined by current and future ANN models of the ventral stream. For instance here we found that synthesis of only few images leads to higher neural response levels that was possible by searching in a relatively large space of natural images (n=640) and complex curved stimuli (n=370) derived from those prior intuitions. This shows that even today’s ANN models – which are clearly not yet perfect (*1, 6*) – already give us new ability to find manifolds of more optimal stimuli for each neural site at a much finer degree of granularity and to discover such stimuli unconstrained by human intuition and difficult to fully describe by human spoken language (see examples in Fig. S1). This is likely to be especially important in mid and later stages of the visual hierarchy (e.g. in V4 and inferior temporal cortex) where the response complexity and larger receptive fields of neurons makes manual search intractable.

In light of these results, what can we now say about the two important critiques of today’s ANN models raised at the outset of this study (understanding and generality)? In our view, the results strongly mitigate both of those critiques, but they do not eliminate them.

On understanding: the ability to use knowledge to gain improved control over things of interest in the world (as we have demonstrated here) is an important test of understanding. However we acknowledge that this is not the only possible view, and many other notions of “understanding” remain to be explored to see if and how these models add value.

On generality: because we found that even today’s ANN models show good generalization to entirely novel images, we believe these results close the door on critiques that argue that current ANN models are extremely narrow in the scope of images they can accurately cover. However, we note that while 54% of the explainable variance in the generalization test was successfully predicted, this is lower than the 89% explainable variance that is found for images that are “closer” to (but not identical to) the mapping images. This not only confirms that these brain models are not yet perfect, but also suggests that a single metric of model similarity to each brain area is insufficient to characterize and distinguish among alternative models (e.g. (*1*)). Instead, multiple similarity tests at different generalization “distances” could be useful, as we can imagine future models that show less decline in successfully predicted variance as one moves from images “near” the training and mapping distributions (typically photographs and naturalistic images) to “far” (e.g. synthetic images like those use here, and others).

From an applications standpoint, the results presented here show how today’s ANN models of the ventral stream can already be used to achieve improved non-invasive, population control (e.g. Fig 4). However, the control results are clearly not yet perfect. For example, in the *one-hot population* control setting we were not able to fully suppress each and every one of the responses of the “off-target” neural sites while keeping the target neural site active (see examples in Figures-3, 4). Post-hoc analysis showed that we could partially anticipate which off-target sites would be most difficult to suppress – they were typically (and not surprisingly) the sites that had high patterns of response similarity with the target site (*r* = 0.49, *p <* 10^*−*4^; correlation between response similarity with the target neural site over naturalistic images and the off-target activity level in the full image one-hot population experiments; n=37 off-target sites). Such results raise very interesting scientific and applied questions of if and when perfect independent control is possible at neuron-level resolution. Are our current limitations on control due to anatomical connectivity that restricts the potential population control, the non-perfect accuracy of the current ANN models of the ventral stream, non-perfect mapping of the model neurons to the individual neural site in the brain, inadequacy of the controller image synthesis algorithm, or some combination of all of these and other factors?

Consider the synthesis algorithm: Intuitively, each particular neural site might be sensitive to many image features, but maybe only to a few that the other neural sites are not sensitive to.

This intuition is consistent with the observation that, using the current ANN model, it was more difficult for our synthesis algorithm to find good controller images in the *One-hot-population* setting than in the *Stretch* setting (the one-hot-population optimization typically took more than twice as many steps to find a synthetic image that is predicted to drive the target neural site response to the same level as in the *Stretch* setting), and visual inspection of the images suggests that the one-hot-population images have fewer identifiable “features” (Figure 5). As the size of the to-be-controlled neural population is increased, it would likely become increasingly difficult to achieve fully independent control, but this is an open experimental question.

Consider the current ANN models: Our data suggest that future improved ANN models are likely to enable even better control. For example, better ANN V4 population predictor models generally produced better one-hot population control of that V4 population (Fig. S5). One thing is clear already – improved ANN models of the ventral visual stream have led to control of high-level neural population that was previously out of reach. With continuing improvement of the fidelity of ANN models of the ventral stream (*1, 25, 26*), the results presented here have only scratched the surface on what is possible with such implemented characterizations of the brain’s neural networks.

## Acknowledgments

We thank Dr. A. Pasupathy for generously providing the complex curvature stimuli, and Kailyn Schmidt for technical support. We also thank Chris Stawarz and the MWorks consortium (https://mworks.github.io) for experimental software support. This research was supported by Intelligence Advanced Research Projects Agency (IARPA), the MIT-IBM Watson AI Lab, US National Eye Institute grants R01-EY014970 (J.J.D.), and Office of Naval Research MURI-114407 (J.J.D).

## Supplementary materials

### Methods

#### Electrophysiological Recordings in Macaques

We sampled and recorded neural sites across the macaque V4 cortex in the left, right, and left hemisphere of three awake, behaving macaques, respectively. In each monkey, we implanted one chronic 96-electrode microelectrode array (Utah array), immediately anterior to the lunate sulcus (LS) and posterior to the inferior occipital sulcus (IOS), with the goal of targeting the central visual representation (<5° eccentricity, contralateral lower visual field). Each array sampled from ~25 mm2 of dorsal V4. On each day, recording sites that were visually-driven as measured by response correlation (*r_pearson_* > 0.8) across split-half trials of a fixed set of 25 out-of-set naturalistic images shown for every recording session (termed, the normalizer image set) were deemed “reliable”.

We do not assume that each V4 electrode was recording only the spikes of a single neuron. Hence we use the term neural “site” throughout the manuscript. But we did require that the spiking responses obtained at each V4 site maintained stability in its image-wise “fingerprint” between the day(s) that the mapping images were tested (i.e. the response data used to build the ANN-driven predictive model of each site, see text) and the days that the Controller images or the complex curvature images were tested (see below). Specifically, to be “stable,” we required an image-wise Pearson correlation of at least 0.8 in its responses to the normalizer set across recording days.

Neural sites that were reliable on the experimental mapping day and the experimental test days, and were stable across all those days, were termed “validated.” All validated sites were included in all presented results. (Note that, to avoid any possible selection biases, this selection of validated sites was done on data that were completely independent from the main experimental result data.) In total, we recorded from 107 validated V4 sites during the ANN-mapping day which included 52, 33 and 22 sites in monkey-M (left hemisphere), monkey-N (right hemisphere), and monkey-S (left hemisphere), respectively. Of these sites, 76 of were validated for the Stretch control experiments (*n_M_* =38, *n_N_* =19, *n_S_*=19) and 57 were validated for the One-hot population control experiments (*n_M_* =38, *n_S_*=19).

To allow meaningful comparisons across recording days and across V4 sites, the raw spiking rate of each site from each recording session was normalized (within just that session) by subtracting its mean response to the 25 normalizer images and then dividing by the standard deviation of its response over those normalizer images (these are the arbitrary units shown as firing rates in Figs. 2A, 3A and 4). The normalizer image set was always randomly interleaved with the main experimental stimulus set(s) run on each day.

Control experiments consisted of three steps. In the first step, we recorded neural responses to our set of naturalistic images that were used to construct the mapping function between the ANN activations and the recorded V4 sites. In a second, offline step, we used these mapping functions (i.e. a predictive model of the neural sites) to synthesize the controller images. Finally in step three, we closed the loop by recording the neural responses to the synthesized images. The time between step 1 and step 3 ranged from several days to 3 weeks.

#### Fixation Task

All images were presented while monkeys fixated a white square dot (0.2°) for 300 ms to initiate a trial. We then presented a sequence of 5 to 7 images, each ON for 100 ms followed by a 100 ms gray blank screen. This was followed by a water reward and an inter-trial interval of 500 ms, followed by the next sequence. Trials were aborted if gaze was not held within ±0.5° of the central fixation dot during any point. To estimate the classical receptive field (cRF) of each neural site, we flashed 1°×1° white squares across the central 8° of the monkeys’ visual field, measured the corresponding neural responses, and then fitted a 2D Gaussian to the data. We defined 1-std as the cRF of each site.

#### Naturalistic Image Set

We used a large set (N=640) of naturalistic images to measure the response of each recorded V4 neural sites and every model V4 neural site to each of these images. Each of these images contained a three-dimensional rendered object instantiated at a random view overlaid on an unrelated natural image background, see (*28*) for details.

#### Complex Curvature Stimuli

We used a set of images consisting of closed shapes constructed by combining concave and convex curves (*12*). These stimuli are constructed by parametrically defining the number and configuration of the convex projections that constituted the shapes. Previous experiments with these shapes showed that curvature and polar angle were quite good at describing the shape tuning (*12*). The number of projections varied between 3 to 5 and the angular separation between projections was in 45° increments. These shapes were previously shown to contain good drivers of V4 neurons of macaque monkeys (*12,14*). The complex curve images were generated using the code generously supplied by the authors of that prior work (http://depts.washington.edu/shapelab/resources/stimsonly.php). The stimuli were presented at the center of the receptive field of the neural sites (detailed below).

#### Cross-Validation Procedure for Evaluating Control Scores

To evaluate the scores from the neural responses to an image set, we divide the neural response repetitions into two, randomly-selected halves. We then compute the mean firing rate of each neural site in response to each image in each half. The mean responses from the first half are used to find the image that produces the highest score (in that half) and the response to that image is then measured in the second half (and this is the measurement used for further analyses). We repeat this procedure 50 times for each neural site (i.e. 50 random half splits). For *Stretch* and *One-hot population* experiments the score functions were the “neural firing rate” and “softmax score” respectively. We compute each score for the synthetic controller images and for the reference images (either the naturalistic or the complex curvature sets, see text). The synthetic “gain” in the control score is calculated as the difference between the synthetic controller score and the reference score, divided by the reference score.

#### V4 encoding model

To use the ANN model to predict each recorded neural site (or neural population), the internal V4-like representation of the model must first be mapped to the specific set of recorded neural sites. The assumptions behind this mapping are discussed elsewhere (*9*), but the key idea is that any good model of a ventral stream area must contain a set of artificial neurons (a.k.a. features) that, together, span the same visual encoding space as the brain’s population of neurons in that area (i.e. the model layer must match the brain area up to a linear mapping). To build this predictive map from model to brain, we started with a specific deep ANN model with locked parameters. Here we used a variant of Alexnet architecture trained on Imagenet (*13*) as we have previously found the feature space at the output of Conv-3 layer of Alexnet to be a good predictor of V4 neural responses (we here refer to this as model “V4”). We used the same training procedure as was described in (*13*), except we did not split the middle convolutional layers between GPUs.

In addition, the input images were transformed using an eccentricity-dependent function that mimics the known spatial sampling properties of the primate retinae. We termed this the “retinae transformation”. We had previously found that training deep convolutional ANN models with retinae-transformed images improves the neural prediction accuracy of V4 neural sites (an increase in explained variance by ~ 5 *−* 10%). The ”retinae transformation” was implemented by a fish-eye transformation that mimics the eccentricity-dependent sampling performed in primate retinae (code available at https://github.com/dicarlolab/retinawarp). All input images to the neural network were preprocessed by randomly cropping followed by applying the fish-eye transformation. Parameters of the fish-eye transformation were tuned to mimic the cones density ratio in fovea at 4° peripheral vision (*29*).

We used the responses of the recorded V4 neural sites in each monkey and the responses of all the model “V4” neurons to build a mapping from model to the recorded population of V4 neural sites (Figure 1). We used a convolutional mapping function that significantly reduces the neural prediction error compared to other methods like principal component regression. Our implementation was a variant of the 2-stage convolutional mapping function proposed in (*30*). The first stage of the mapping function consists of a learnable spatial mask (*W_s_*) that is parameterized separately for each neural site (*n*). The second stage consists of a mixing pointwise convolution (*W_d_*) that computes a weighted sum of all feature maps at a particular layer of the ANN model (Conv3 layer in our case). The final output is then averaged over all spatial locations to form a scalar prediction of the neural response. Parameters are jointly optimized to minimize the prediction error *L_e_* on the training set regularized by combination of *L*_2_ and smoothing Laplacian losses *L_laplace_* (defined below).

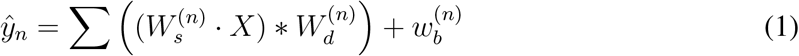

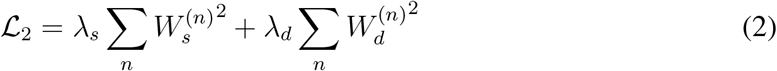

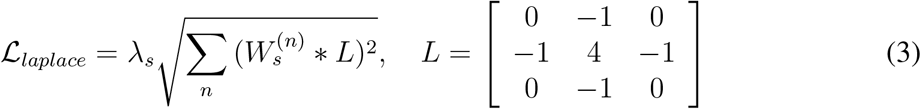

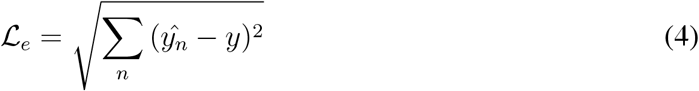

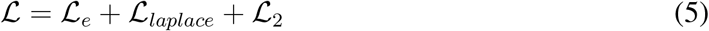

We evaluated our model using 2-fold cross-validation and observed that ~89% of the explainable variance could be explained with our model in three monkeys (*EV_M_* = 92%, *EV_N_* = 92%, *EV_S_* = 80%). The addition of the retinae transformation together with the convolutional mapping function increased the explained variance by ~13% over the naive principal component regression applied on features from the model trained without the retinae transformation (two monkeys: *EV_M_* = 75%, *EV_N_* = 80%, *EV_S_* = 73%). For constructing the final mapping function, adopted for image synthesis, we optimized the mapping function parameters on 90% of the data, selected randomly.

The resulting predictive model of V4 (ANN features plus linear mapping) is referred to as the *mapped v4 encoding model* and, by construction, it produces the same number of artificial V4 “neurons” as the number of recorded V4 neural sites (52, 33, and 22 neural sites in monkeys M, N and S respectively).

#### Synthesized “Controller” Images

The “response” of artificial neuron in the *mapped V4 encoding model* (above) is a differentiable function of the pixel values *f* : *I^w×h×c^ →* ℝ*^n^* that enables us to use the model to analyze the sensitivity of neurons to patterns in the pixels space. We formulate the synthesis operation as an optimization procedure during which images are synthesized to control the neural firing patterns in the following two settings:

1. **Stretch:** We synthesize controller images that attempt to push each individual V4 neural site into its maximal activity state. To do so, we iteratively change the pixel values in the direction of the gradient that maximizes the firing rate of the corresponding model V4 neural site. We repeated the procedure for each neural site using five different random starting images, thereby generating five “stretch” controller images for each V4 neural site.
2. **One Hot Population:** Similar to “Stretch” scenario, except that here we chose the optimization to change the pixel values in a way that (i) attempts to maximize firing rate of the target V4 neural site, and (ii) attempts to maximally suppress the firing rates of all other recorded V4 neural sites. We formalize the *One-hot population* goal in the following objective function that we then aim to maximize during the image synthesis procedure:

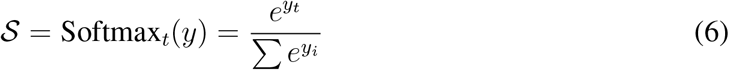

where *t* is the index of the target neural site, and *y_i_* is the response of the model V4 neuron *i* to the synthetic image.

For each optimization run, we start from an image that consists of random pixel values drawn from a standard Normal distribution and optimize the objective function for a prespecified number of steps using gradient ascend algorithm (steps=700). During the experiments, monkeys are required to fixate within a 1° circle at the center of the screen. This introduces an uncertainty on the exact gaze location. For this reason, images are synthesized to be robust to small translations of maximum 0.5°. At every iteration, we translate the image in random directions (i.e. jittering) with a maximum translation length of 0.5° in each direction, thereby, generating images that are predicted to elicit similarly high scores regardless of the translations within the range.

#### Contrast Energy

It has been shown that neurons in area V4 respond more strongly to higher contrast stimuli (*31*). To ask if contrast energy (CE) was the main factor in “stretching” the V4 neural firing rates, we computed the contrast energy within the receptive field of the neural sites for all the synthetic and the classic V4 stimuli. Contrast energy was calculated as the ratio between the maximum and background luminances. For all images, the average luminance was used as the background value. Because the synthetic images consisted of complex visual patterns, we also computed the contrast energy using an alternative method based on spectral energy within the receptive field. We calculated the average power in the cRF in the frequency range of 1-30 cycles/degree. We ensured that for all tested neural sites, CE within the cRF for synthetic *Stretch Controller* images were less than or equal to the classic, complex curvature V4 stimuli (Supp Figure-S4).

**Figure S1:**
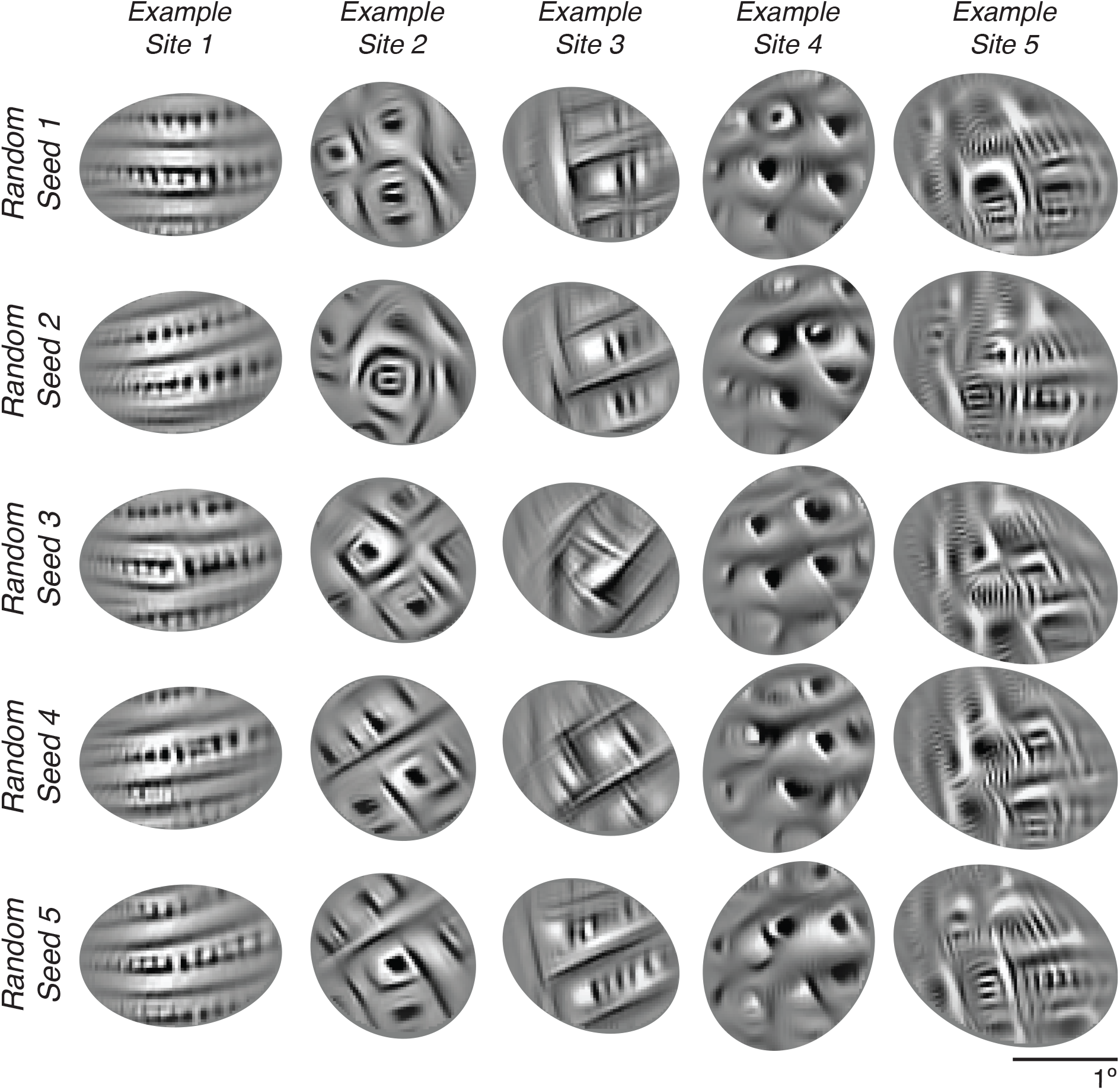
Stretch synthetic controller images for five example V4 neural target sites (Example sites 1-5). Each row displays images generated using the same random starting image, but optimized for each target site. Note the perceptual similarity of the controller images synthesized for each site and the dissimilarity between the controller images across sites.

**Figure S2:**
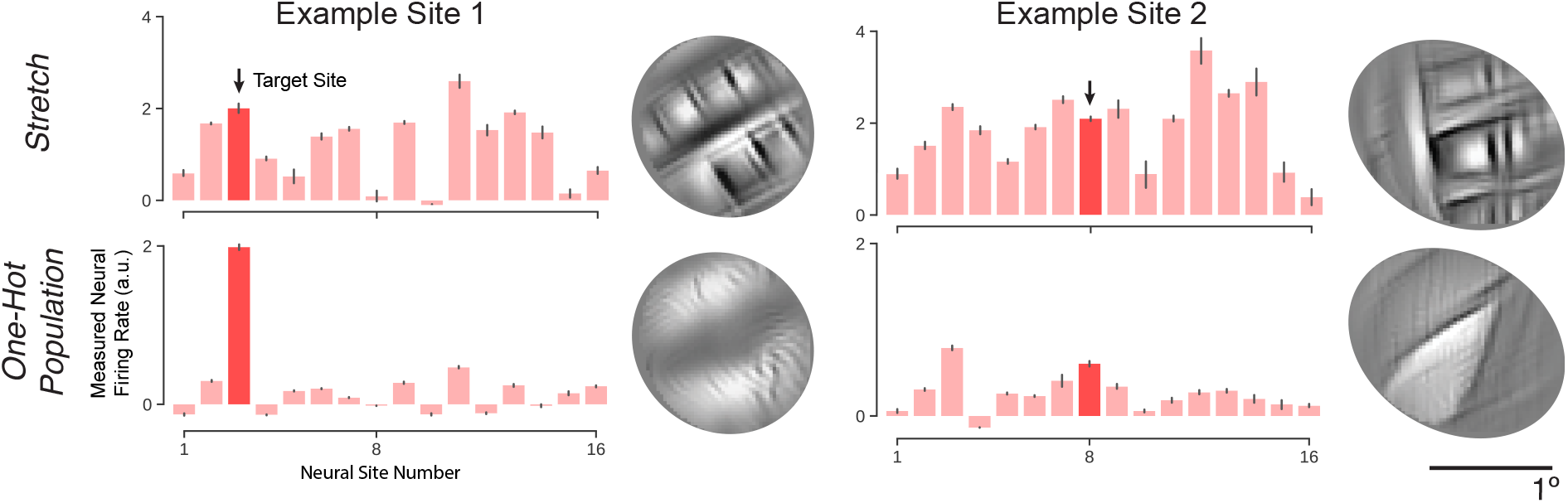
Comparison of population response in *Stretch* and *One-hot Population* settings. Population responses in *Stretch* and *One-hot Population* settings are demonstrated for two example neural sites. One-hot population images were generated with an objective function including 16 neural sites with highly overlapping receptive fields. Compared to the *Stretch* controller images, the *one-hot-population* images have fewer identifiable “features”. The displayed images were synthesized using the same initial random image.

**Figure S3:**
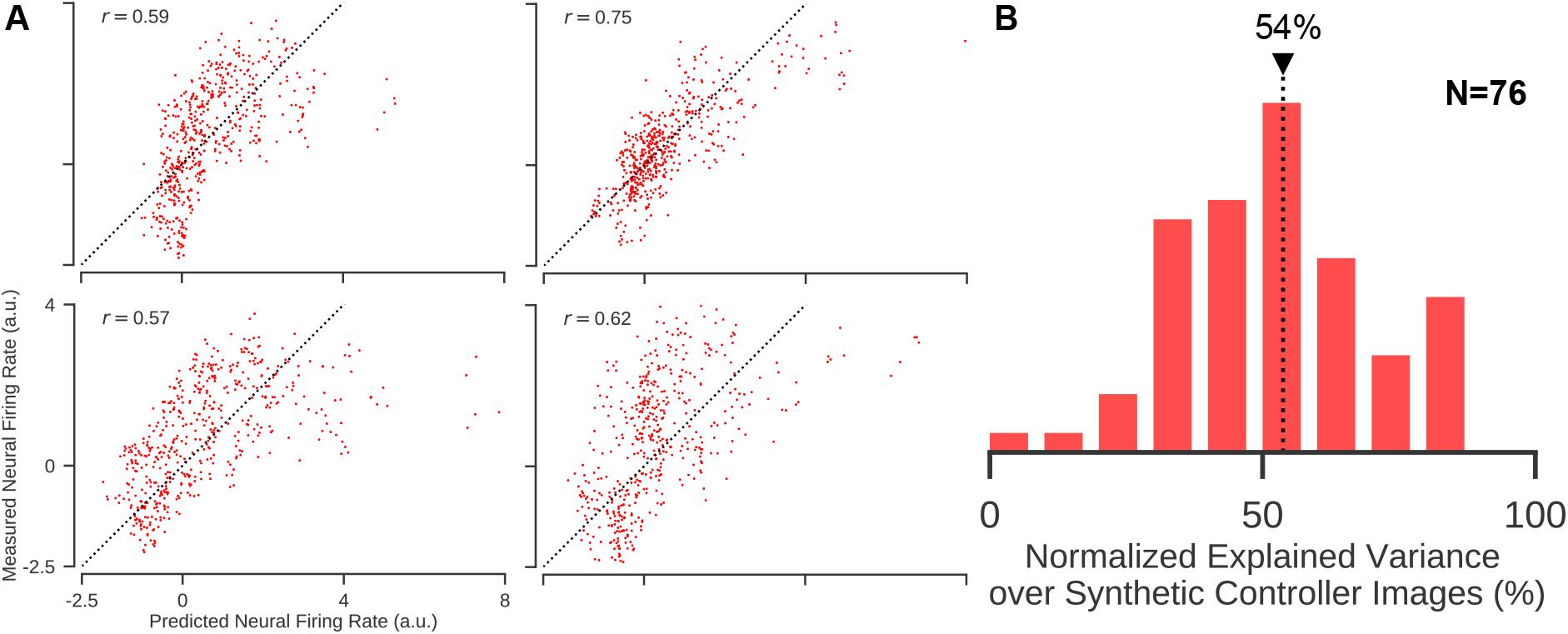
Predictability of synthetic controller images. A) Scatter plots of predicted and measured V4 neural responses to synthetic controller images for four example neural sites. For most target neural sites, the predicted and measured neural responses were significantly correlated. Each dot represents the prediction and average measured response to a single image. B) The model accounted for 54% (median across all tested neural sites in three monkeys; *N* =76) of the explainable variance.

**Figure S4:**
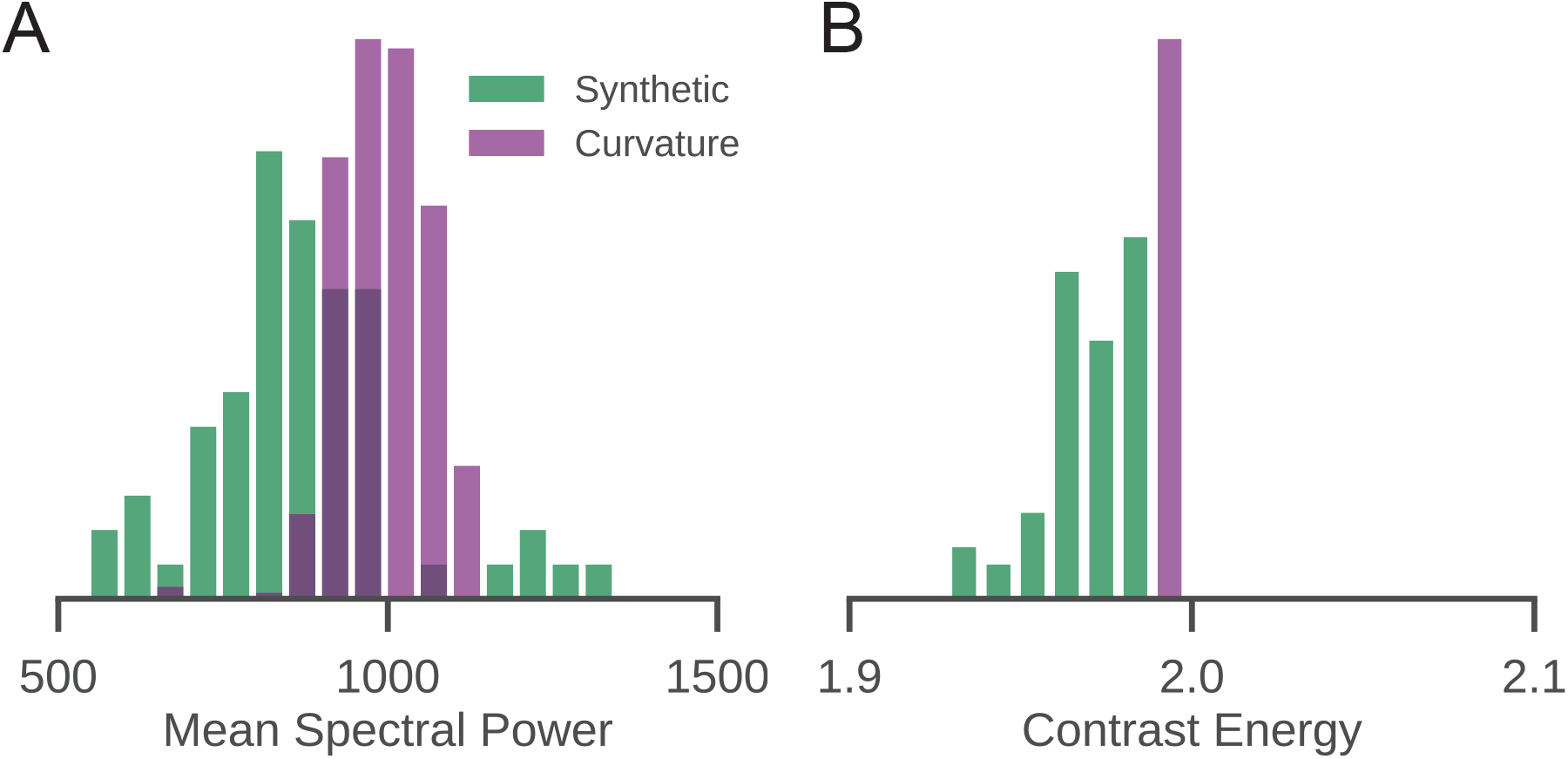
Comparison of contrast energy between synthetic and curvature images. A) Distribution of the mean spectral power within target neural sites’ classic receptive fields for “Stretch” controller (green) and complex curvature (purple) images. Spectral power was computed using 2-D FFT transformation and summed in the frequency range of 1-30 cycles/degree. B) Distribution of contrast energy within target neural sites’ classic receptive fields for “Stretch” controller and complex curvature images.

**Figure S5:**
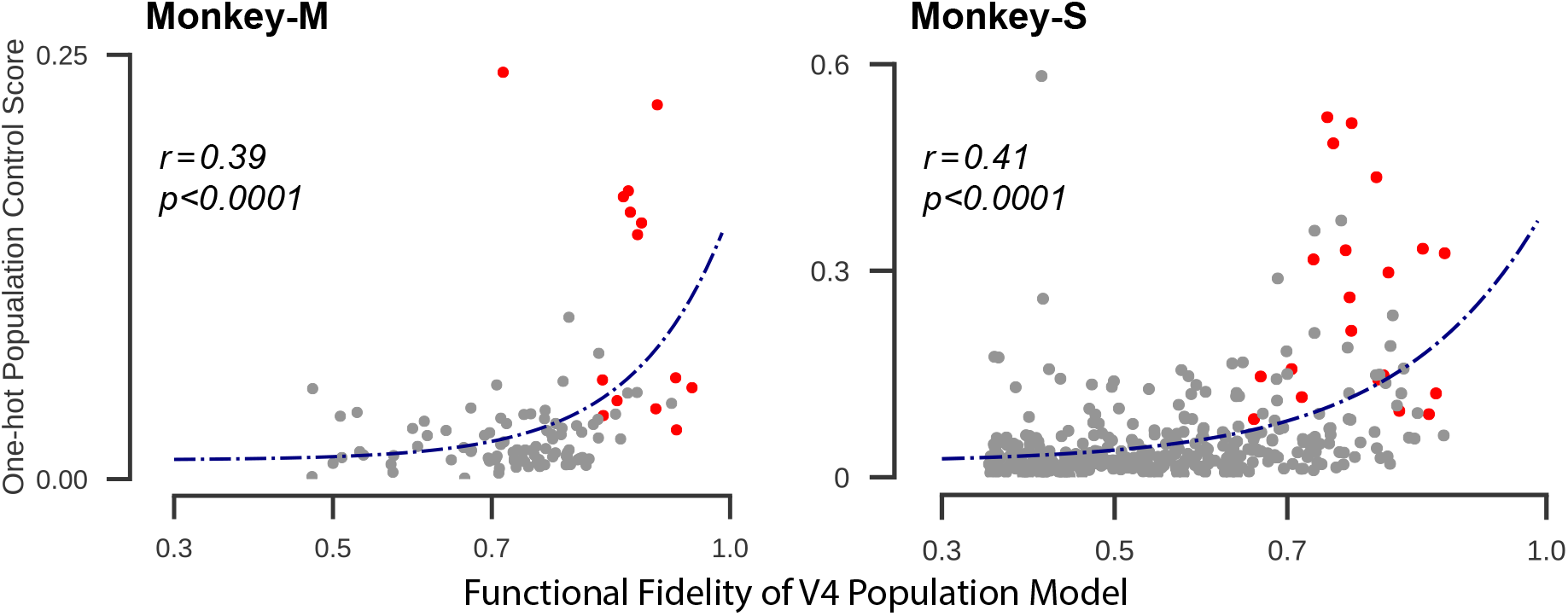
Higher functional fidelity models increase the ability to control neural responses. We evaluated the one-hot population control score for each target neural site in each monkey subject for a range of possible models with different prediction accuracy levels. In each monkey session, the functional fidelity of a V4 population model (measured by the mean of: 1) explained variance of target neural site and 2) the mean of the explained variance for all the off-target sites) was plotted against the one-hot population control score achieved with that population model. We found that these were significantly correlated as assessed by Spearman rank order correlation, shown on each panel. For this analysis, for each target neural site, we included not only the original tests, but also tests in which we swapped the predictive model of the target neural site with the model of randomly-chosen off-target site (we do this ”mismatch” test because it is an example of what would have happened in the experiment if the synthesis algorithm had been given the wrong models – it would have produced OHP control stimuli that we already tested – so we can compute the resulting control score without doing new recording experiments). We simply assessed the functional fidelity of V4 population model using the mismatched models and the population control score achieved using the new population model’s synthetic control images (again, from population responses to images that we had already tested). Red dots correspond to cases where the target neural site’s model and responses were matched (i.e. results of the original OHP tests, see text), and gray dots correspond to the cases where they were mismatched. Dark blue line shows an exponential function fitted to the data points, highlighting the tendency for higher model fidelity to support better control.

